# Multi-organ structural homogeneity of amyloid fibrils in ATTRv-T60A amyloidosis patients, revealed by Cryo-EM

**DOI:** 10.1101/2024.05.14.594218

**Authors:** Maria del Carmen Fernandez-Ramirez, Binh A. Nguyen, Preeti Singh, Shumaila Afrin, Bret Evers, Parker Basset, Lanie Wang, Maja Pękała, Yasmin Ahmed, Virender Singh, Jacob Canepa, Aleksandra Wosztyl, Yang Li, Lorena Saelices

## Abstract

ATTR amyloidosis is a degenerative disorder characterized by the systemic deposition of the protein transthyretin. These amyloid aggregates of transthyretin (ATTR) can deposit in different parts of the body causing diverse clinical manifestations. Our laboratory aims to investigate a potential relationship between the different genotypes, organ of deposition, clinical phenotypes, and the structure of ATTR fibrils. Using cryo-electron microscopy, we have recently described how the neuropathic related mutations ATTRv-I84S and ATTRv-V122Δ can drive structural polymorphism in *ex vivo* fibrils. Here we question whether the mutation ATTRv-T60A, that commonly triggers cardiac and neuropathic symptoms, has a similar effect. To address this question, we extracted and determined the structure of ATTR-T60A fibrils from multiple organs (heart, thyroid, kidney, and liver) from the same patient and from the heart of two additional patients. We have found a consistent conformation among all the fibril structures, acquiring the “closed-gate morphology” previously found in ATTRwt and others ATTRv related to cardiac or mixed manifestations. The closed-gate morphology is composed by two segments of the protein that interact together forming a polar channel, where the residues glycine 57 to isoleucine 68 act as a gate of the polar cavity. Our study indicates that ATTR-T60A fibrils present in peripheral organs adopt the same structural conformation in all patients, regardless of the organ of deposition.

## Introduction

ATTR amyloidosis is a degenerative disease with diverse clinical manifestations, characterized by the pathologic deposition of amyloidogenic transthyretin (ATTR). This deposition stems from mutations in hereditary ATTR (ATTRv) amyloidosis, or unknown aging-related factors in wild-type ATTR (ATTRwt) amyloidosis. Contrary to neurodegenerative diseases such as Alzheimer’s or Parkinson’s, the amyloid deposition of ATTR occurs far from the secreting organ, which is mainly the liver, and affects multiple tissues of the body. This systemic deposition results in a variety of symptoms, with the most common presentations being polyneuropathy, cardiomyopathy, or a mixed phenotype (Hazenberg, 2013). The molecular basis of this phenotypic variability is unknown.

In the past years, our laboratory has set out to investigate whether the diverse clinical presentation of ATTR amyloidosis is associated with the structural conformation of ATTR fibrils. We started this quest after recent cryo-EM studies of brain fibrils from patients of neurodegenerative diseases revealed that each disease is associated with specific fibril conformation(s), suggesting a connection between fibril structure and pathology (Arseni et al., 2022; Fitzpatrick et al., 2017; Scheres et al., 2020; Schweighauser et al., 2020; Yang et al., 2022). Our data, together with the studies from others (Iakovleva et al., 2021; Schmidt et al., 2019; Steinebrei et al., 2023; Steinebrei et al., 2022), revealed that all ATTR fibrils share a common core composed by two TTR fragments with a polar pocket formed by the C-terminal fragment. The two fragments interact by a three-sided interface of interdigitated steric zippers (B. A. Nguyen et al., 2024). Despite the high structural homogeneity of ATTR fibrils, we and others have found structural variations, or *polymorphism*, in ATTR fibrils.

ATTR fibril polymorphism was found at two levels: the number of protofilaments and the conformation of the polar pocket. With the exception of fibrils from an ATTRv-V122Δ patient (Yasmin et al., 2024), all cardiac ATTR fibrils are found to have one single protofilament (Binh An Nguyen et al., 2024; B. A. Nguyen et al., 2024; Schmidt et al., 2019; Steinebrei et al., 2023; Steinebrei et al., 2022). The exceptions, ATTRv-V122Δ cardiac fibrils from a neuropathic patient, contained one or two protofilaments; with single fibrils being the most represented population (78%). This multiplicity of protofilaments was also observed in one other organ, the eye. Around 80% of the fibrils extracted from the vitreous body of the eye of an ATTRv-V30M patient contained multiple protofilaments (singles, doubles, and triples) (Iakovleva et al., 2021). As for the conformation of the polar pocket, most cardiac ATTR fibrils studied by cryo-EM acquires what we called a “closed-gate” conformation, in where the segment from glycine 57 to glycine 67 closes the polar pocket forming a channel (B. A. Nguyen et al., 2024; Schmidt et al., 2019; Steinebrei et al., 2023; Steinebrei et al., 2022). In contrast, this segment can adopt multiple conformations in cardiac ATTRv-I84S fibrils (B. A. Nguyen et al., 2024) and ATTRv-V30M fibrils from the eye (Iakovleva et al., 2021). The ATTRv-I84S and ATTRv-V122Δ patients included in these studies manifested ATTRv-related polyneuropathy (ATTR-PN). The rest of patients studied by cryo-EM so far, both with ATTRwt and ATTRv amyloidosis, manifested ATTR-related cardiomyopathy (ATTR-CA) or a mixed phenotype (Adams et al., 2021). This observation opens the question of whether the ATTR-CA fibrils could be associated with a more homogeneous structural landscape whereas ATTR-PN fibrils could be more polymorphic. However, these structural differences could also -or rather- be dependent on mutation, affected organ, or other factors.

Here we question whether the mutation ATTRv-T60A drives fibril polymorphism and if its deposition in different organs can cause different morphologies. The mutation of threonine to alanine in position 60 is one of the over 140 single-point pathological mutations in the *ttr* gene. Many of these residue substitutions are thought to destabilize the native tetrameric form of transthyretin, releasing amyloidogenic monomers that partially unfold and aggregate (Hammarström et al., 2002; Ruberg & Berk, 2012). The mutation T60A causes an increase of flexibility in one of the outside loops in the native state (Cendron et al., 2009) and is thermodynamically less stable than the wild-type transthyretin (Sekijima et al., 2005). It originated in north-west of Ireland (County Donegal) and is present in 1% of its population. Then, it spread to the rest of the United Kingdom, the Appalachian region of the United States, Europe, and Japan (Barker & Judge, 2022). ATTRv-T60A amyloidosis presents as a mixed phenotype of neuropathy and cardiomyopathy, with a dominant cardiac phenotype that is a major determinant of its poor prognosis (Hewitt et al., 2020; Sattianayagam et al., 2012).

In this manuscript, we disclose six structures of ATTRv-T60A fibrils collected from three patients, and from multiple organs, including the heart, liver, kidney, and thyroid from one individual, and the hearts from two additional individuals. All these structures acquired the aforementioned “closed-gate” conformation, with the segment from glycine 57 to glycine 67 closing the gate of the polar pocket into a channel. Our study provides the first multi-organ description of fibril structures from the same ATTR amyloidosis patient, and suggests that ATTR-T60A fibrils adopt a consistent structural conformation in all patients, regardless of the organ of deposition. These findings contribute to the understanding of the biopathology of ATTR aggregation by providing evidence that the local environment of the deposition site may not necessarily drive fibril polymorphism.

## Results

### ATTR-T60A fibrils from the heart of three different patients share a common fold

ATTR fibrils were extracted from the heart of three individuals that carried the *ttr* mutation T60A in heterozygosis. Heart samples from the patients 1 and 2 were freshly frozen while that from the patient 3 came from lyophilized fibril extracts obtained by the team of Dr. Benson in Indiana University, following a previously described protocol (Liepnieks & Benson, 2007). We used a water-based extraction protocol previously described to further purify fibrils from fresh tissue or lyophilized material (Binh An Nguyen et al., 2024; B. A. Nguyen et al., 2024). The different sample elutions were visualized by electron microscopy **(Supplementary Fig. 1a)** and immuno-dot blot in order to confirm the success of the extraction, optimize the sample preparation when needed, and select the one with higher quality for the cryo-EM grid preparation. We plunge-freeze our samples by using a Vitrobot, and did several rounds of screening and optimization of our grids until we achieve high-quality preparation for the data collection.

We started the data processing with rounds of 2D classification. At this step, we identified two types of fibrils as we previously described (B. A. Nguyen et al., 2024). Most 2D classes corresponded to twisted fibrils, while other classes showed stacked layers without apparent twist **(Supplementary Fig. 1b)**. These untwisted classes were removed for the next rounds of 2D classification. Visually, only one twisted morphology was distinguishable in the three patients. The twisted 2D classes from this morphology were stitched together to obtain crossover distances for helical reconstruction **(Supplementary Fig. 1b)**.

For the three patients, 3D classification of the twisted fibrils resulted in only one distinguishable class. The final density maps for the fibrils of the three patients have a resolution of 3.18 Å for patient 1, 3.4 Å for patient 2, and 4.64 Å for patient 3 **(Figure 1, Supplementary Fig. 2a)**, and their crossover distances are 700 Å, 677 Å, and ∼690 Å, respectively. For patients 1 and 2, their twist angles are –1.26 and -1.27, and the rise between layers are 4.9 Å and 4. 78 Å. These values could not be calculated for patient 3 (**Supplementary Table 1)**.

**Figure 1.**
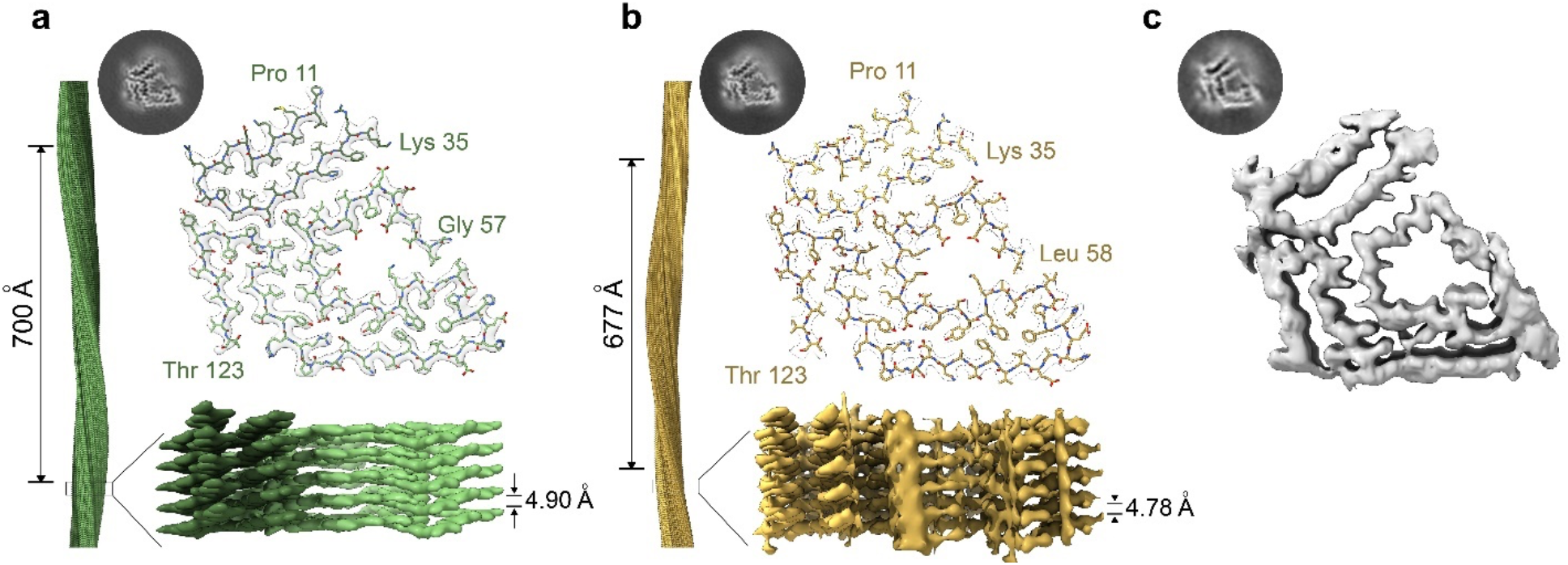
Cryo-EM structures of cardiac fibrils from three different ATTRv-T60A patients. 3D class and models of the structures resolved from the fibrils of **a)** Patient 1 and **b)** Patient 2. Although different interlayer distance and crossover were calculated, we can see they share the same morphology composed by two fragments of transthyretin: N-terminal (proline 11 to lysine 35) and C terminal (Threonine 123 to glycine 57/Leucine 58). **c)** The 3D class obtained from patient 3 and the morphology distinguished in the density map indicate that these fibrils acquired the same morphology previously described for patient 1 and 2. The density map obtained from this patient had not resolution enough to construct the model.

The built models (Patient 1 and Patient 2) revealed that the ATTRv-T60A fibrils deposited in the heart of these three different patients share a common morphology. The core is composed of two fragments of the transthyretin protein: an N terminal fragment that includes residues from proline 11 to arginine 34 (in patient 1) or to lysine 35 (in patient 2), and a C terminal fragment that includes residues from glycine 57 (in patient 1) or leucine 58 (in patient 2) to residue threonine 123 **(Figure 1 a-b)**. Some of these β-strands acquired contribute to the formation of steric zippers, composed mainly of hydrophobic residues **(Figure 1 a-b** and **Supplementary Fig. 2d)**. The analysis of their solvation energies revealed that this configuration creates three areas that are estimated to be energetically stable: two pockets formed by hydrophobic residues and the interface that connects the N- and C-fragments together **(Supplementary Fig. 2c)**. These areas may be responsible for their estimated stability, which are significantly higher than the rest of amyloid structures analyzed so far (M. R. Sawaya et al., 2021). The ionizable or hydrophilic residues are exposed to the outside of the fibril, as expected, and they are also located facing a cavity in where the segment from glycine 57 to glycine 67 (previously referred to as the gate) closes the polar pocket forming a channel.

### ATTR-T60A amyloid fibrils were deposited in the kidney, liver, and thyroid of patient 3

The evaluation of other tissues from patient 3 confirms amyloid deposition. We performed a histological analysis of the thyroid, kidney, and liver of patient 3, who underwent a kidney-liver transplantation, to confirm or rule out the presence of ATTR amyloid deposits. We selected different anatomical regions of the organ, and we stained the subsequent slides with hematoxylin and eosin, Congo red, and Thioflavin S. The fat found in the different organs presented amyloid deposition **(Figure 2**) while the parenchyma did not show significant amyloid content. This fat was localized surrounding the thyroid and the kidney, while in the liver was attached to the round ligament. We also detected amyloid deposition in the fatty areas of the nerves of the kidney and the Gibson capsule that surrounds the liver. Slices from the parenchyma of the thyroid, show well defined and limited areas with amyloid aggregates attached to the vascular wall **(Figure 2)**.

**Figure 2.**
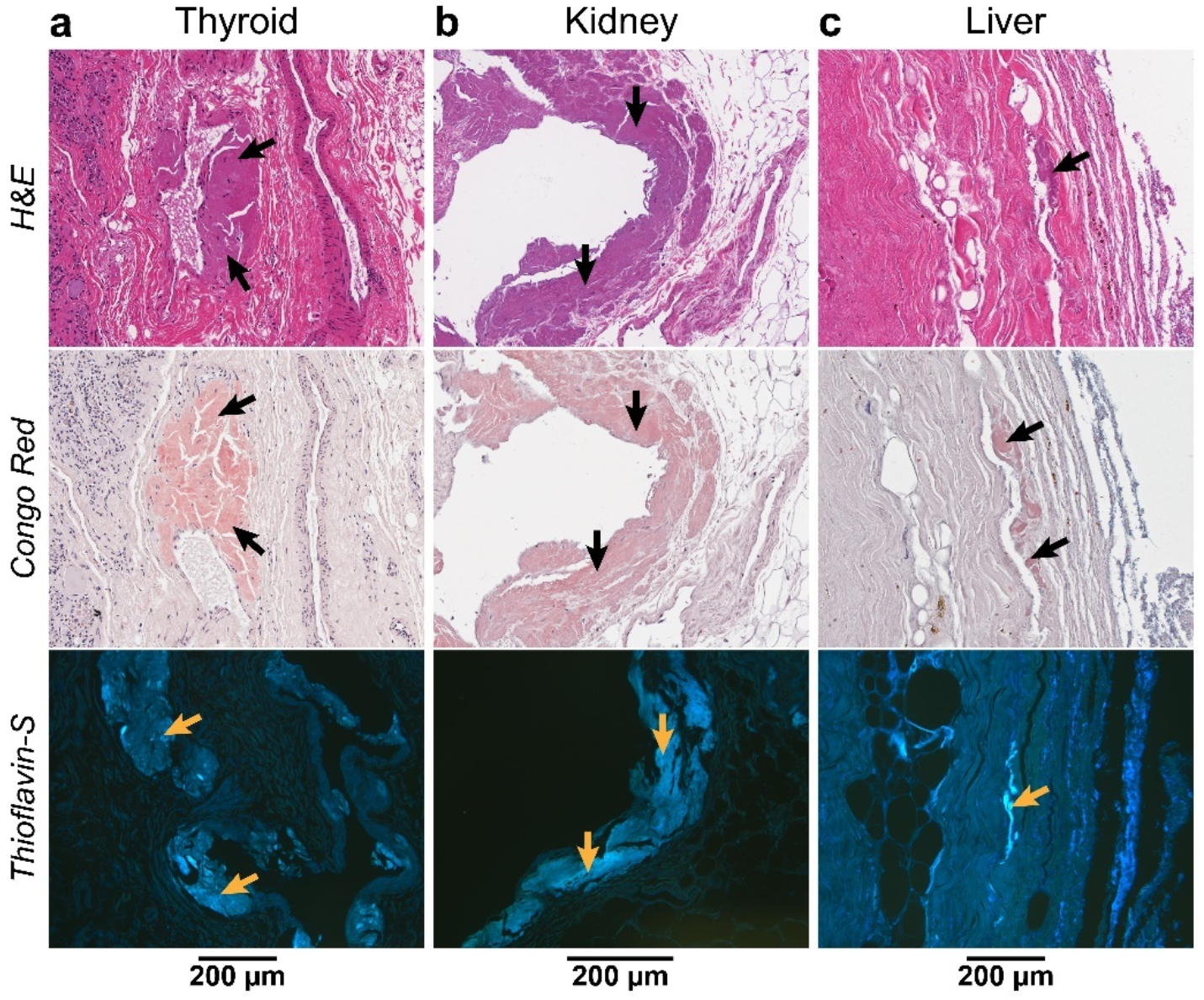
Identification of amyloid deposits by histological staining of the non-cardiac organs from Patient 3. General cellular staining with hematoxylin and eosin (top) and specific amyloid staining with Congo red (middle) and Thioflavin S, Th-S, (bottom) was done for every tissue. **a)** Congo red stained amyloid deposition closed to the blood walls and Th-S showed the deposits found in the adipose tissue of thyroid. **b)** Slices corresponding to the same areas were stained with the different dyes. The amyloid deposits found in adipose tissue are indicated. **c)** Staining of a cross-section slice of the liver show the amyloid-specific staining of the Glisson capsule.

We extracted ATTR fibrils from the three organs following the protocol previously used for the cardiac samples. We selected the fatty areas surrounding the different organs because of their amyloid content shown by histology **(Figure 2)**. As control, we included the parenchyma of the liver. We confirmed the presence of transthyretin in the resulting elutions by immuno-dot blotting. We then visualized the extracted ATTR fibrils by transmission electron microscopy (TEM) from all the elutions, excepting those from the parenchyma than only had few isolated fibrils. All of them visually showed a similar morphology than those found in the cardiac tissue.

### ATTRv-T60A fibrils from other organs are not structurally polymorphic

We observed that the ATTRv-T60A fibrils deposited in the hearts of three distinct patients exhibit the same “closed-gate” conformation, consistent with cardiac ATTR fibrils observed in previous patient studies (Binh An Nguyen et al., 2024; B. A. Nguyen et al., 2024; Schmidt et al., 2019; Steinebrei et al., 2023; Steinebrei et al., 2022). Given that patient 3 also displayed amyloid deposition in other organs, we investigated whether there is an organ-dependent structural variability in ATTRv-T60A fibrils.

We first prepared the samples containing fibrils extracted from thyroid, kidney, and liver for cryo-EM data collection and processing **(Supplementary Fig. 2)**. Similar to the previous cardiac ATTRv-T60A fibrils, we found both straight and curvy classes morphologies in the 2D classification from the data of kidney, liver, and thyroid fibrils. The stitching of the curvy classes reconstructed show only one morphology that generated one 3D class, shared among them and with those from cardiac tissue **(Supplementary Fig. 3)**. In the case of the fibrils obtained from the kidney, we noticed additional annular elements of ∼3.5 nm in diameter, attached to the fibrils **(Supplementary Fig. 3)**. They appeared in the first rounds of 2D classification and then disappeared progressively. We have not yet identified these annular elements.

The 3D classification yielded density maps of the ATTRv-T60A fibrils across different organs, with a resolution of XX Å for those originating from the thyroid, XX Å for those from the kidney, and XX Å for those from the liver. These fibrils display crossover and twist parameters of 701 Å and –1.23, 755 Å and –1.17, and 707 Å and –1.25, respectively. Additionally, the inter-layer distances are measured at 4.79 Å, 4.91 Å, and 4.91 Å for thyroid, kidney, and liver fibrils, respectively. Utilizing these density maps, we constructed three additional models showcasing the “close gate” morphology. These models delineate the N-terminal fragment of TTR spanning from proline 11 to lysine 35 and the C-terminal fragment encompassing residues from glycine 57 to threonine 123, forming the polar cavity sealed by residues from glycine 57 to glycine 67. From these refined structures derived from ATTRv-T60A fibrils across thyroid, kidney, and liver, and thyroid tissues of the same patient, we infer that the morphological characteristics of these fibrils, bearing the T60A mutation, remain consistent irrespective of the organ of deposition.

### Molecular characterization of the ATTRv-T60A fibrils analyzed in this study

The six structures obtained in this study correspond to amyloid fibrils extracted from the hearts of three individuals, along with fibrils sourced from the thyroid, kidney, and liver of one of these patients. Despite their diverse origins (whether from patients or organs), they display a consistent amyloid conformation. In light of this observation, we extended our analysis with additional comparative characterization to identify shared features or potential differences among them.

Mass spectrometry analysis of the fibril extracts provides valuable insights into their composition. Several residues within the transthyretin sequence were not incorporated into the cryo-EM models due to the absence of corresponding density. However, mass spectrometry analysis of the fibrils revealed the full-length sequence of transthyretin, suggesting that these missing residues are not lacking but rather remain unfolded outside the fibril structure. Additionally, we detected the presence of both wild-type and T60A mutant transthyretin, consistent with the heterozygous status of the patients. Analysis of tryptic peptides encompassing residue 60 provided qualitative insights into the abundance of wild-type and mutated transthyretin within the fibrils. We found an approximate wild-type to mutant ratio of 50:50 in the hearts of patients 1 and 2, and 75:25 in the heart of patient 3. For the patient 3, the ratio for the thyroid was 75:25, 50:50 for kidney, and 40:60 for the liver. We notice a higher wild-type prevalence in those tissues from patient 3 that remained in the patient after the combined kidney-liver transplantation.

Given the association proposed between the presence of wild-type transthyretin and fragmented C-terminal transthyretin in amyloid deposits (Ihse et al., 2011), we examined the prevalence of C-terminal fragments in the fibrils included in this study. Western blot of the ex-vivo ATTR fibrils using an antibody against the fragment from residue 50 to 127 revealed the presence of the full-length monomeric transthyretin, with the typical band corresponding to ∼15 kDa and an additional smaller band of ∼8-10 kDa **(Figure 4a)**. In contrast, type B fibrils are only composed of full-length TTR and is commonly related to the ATTRv-V30M variant (Ihse et al., 2011). The intensity ratio between the ∼15 kDa band and the ∼8-10 kDa band does not significantly vary among patients nor organs **(Figure 4b)**, suggesting a stable prevalence of cleaved transthyretin within the fibrils among ATTRv-T60A cases.

We also analyzed the seeding capacity of the ATTRv-T60A fibrils included in this study using recombinant monomeric transthyretin, or MTTR **(Figure 4 c,d)**. Various cardiac ATTR fibrils, including those carrying the mutation ATTRv-T60A, have been reported to trigger the aggregation of the engineered transthyretin variant “MTTR” (F87M/L100M) at physiological pH (Saelices et al., 2018). Under these conditions, MTTR remains in a monomeric state, partially unfolded, and thus poised for amyloid formation (Jiang et al., 2001). Using this engineered variant bypasses the dissociation of the tetramer, which is typically the rate-limiting step for aggregation (Colon & Kelly, 1992). We generated fibril seeds from our fibril extracts by gentle purification with detergent and sonication for fragmentation, as previously described (Saelices et al., 2018). The amyloid formation of MTTR in the presence of these seeds was tracked by monitoring thioflavin T fluorescence over 80 hours. With the exception of the reaction with thyroid seeds, all seeding curves exhibited the characteristic sigmoidal shape of amyloid fibril formation (Xue et al., 2008). The kinetics of all samples showed a lag time ranging from 15 to 26 hours **(Figure 4c)**. We validated these results by visualizing the presence of seeded fibrils by transmission electron microscopy and negative staining **(Figure 4d)**, and the presence of MTTR in the insoluble fraction post-reaction using immuno-dot blotting.

Taken together, our findings indicate that ATTRv-T60A fibrils consist of both wild-type and mutated transthyretin, with the presence of the variant significantly reduced in non-cardiac tissues. These fibrils are classified as type A, composed of both full-length and cleaved C-terminal transthyretin. Except for thyroid fibrils, they all retain the ability to induce the aggregation of recombinant transthyretin *in vitro*, irrespective of the patient or organ of origin.

## DISCUSSION

ATTR amyloidosis is a disease with diverse clinical manifestations that involves the pathological aggregation of transthyretin. Depending on the primary manifestation of the symptoms, ATTR amyloidosis can be classified in three main groups: ATTR-related cardiomyopathy, ATTR-related peripheral neuropathy, and ATTR amyloidosis patients with a mixed phenotype. We hypothesize that this phenotypic variability could be associated with the structural conformation of amyloid fibrils. Here we determined the structures of fibrils obtained from the heart of three ATTRv-T60A amyloidosis patients, and from the thyroid, kidney, and liver of one of those three patients. These patients manifested a mixed phenotype consisting of both cardiomyopathy and polyneuropathy.

### Mixed phenotypes in ATTR amyloidosis may be associated with structural fibril homogeneity

Together with our previous cryo-EM studies of ATTR fibrils, our findings suggest that the polymorphism of ATTR fibrils may be associated with neuropathic phenotypes, rather than cardiomyopathic or mixed phenotypes. ATTRv fibrils were found polymorphic in ATTRv-I84S and ATTRv-V122Δ amyloidosis patients with neuropathy (B. A. Nguyen et al., 2024; Yasmin et al., 2024). In contrast, ATTRwt and ATTRv fibrils from amyloidosis patients with cardiomyopathy or mixed phenotypes were found structurally consistent (Iakovleva et al., 2021; Binh An Nguyen et al., 2024; Schmidt et al., 2019; Steinebrei et al., 2023; Steinebrei et al., 2022). Since the previous structural studies of ATTRv fibrils were constrained by the inclusion of only one patient for each genotype, the ability to establish a direct link between the structure of ATTRv fibrils and specific genotypes is restricted. Here we provide a comprehensive structural study of amyloid fibrils from ATTRv-T60A amyloidosis patients, by including three independent patients, and multiple organs. We observe that in all three cases, the structures of ATTRv-T60A fibrils share the same conformation **(Figure 1)**, corresponding to the close-gate fold observed in ATTRwt and other ATTRv patients with cardiomyopathy and mixed phenotypes. Our findings hint at a potential relationship between fibril structural homogeneity and mixed phenotypes in ATTR amylodiosis patients.

### The structure of ATTRv-T60A fibrils does not depend on the organ of deposition

Previous studies suggest that ATTRv fibril polymorphism may be organ specific (Iakovleva et al., 2021; Schmidt et al., 2019). These studies describe the presence of fibril polymorphs in the vitreous humor of an ATTRv-V30M patient different from the structure found in the heart of a second patient carrying the same mutation. The structural differences in fibrils from the heart and the eye could be driven by at least three factors: the local environment of the deposition site, the local environment of the media in where transthyretin circulates, or protein changes made before secretion by its source (this is, the liver or the retina epithelium, respectively). The importance of the local environment in an amyloid protein to acquire different fibril morphologies has already been reported for tau (Lövestam et al., 2022), modulating the first structural rearrangement of the monomer prior to aggregation (Fernández-Ramírez et al., 2023). These structural rearrangements could occur while the protein is in circulation or at the deposition site. To assess whether the local environment of the deposition site can drive the formation of fibril polymorphs, we determined the structures of ATTRv-T60A fibrils from different organs (thyroid, kidney, liver, and the heart) from the same patient **(Figure 1 and 3)**. We chose these organs because the source of transthyretin for all these peripheral tissues is the same, the liver. The absence of polymorphism in the resulting ATTR structures suggests that the fibril differences observed in ATTRv-V30M may be influenced by factors other than the local environment of the deposition site, for example, the local environment of the media in where transthyretin circulates prior to tissue infiltration (vitreous humor in the eye vs blood in the heart), or changes made before secretion by its source (retina epithelium for the eye vs liver for the heart).

**Figure 3.**
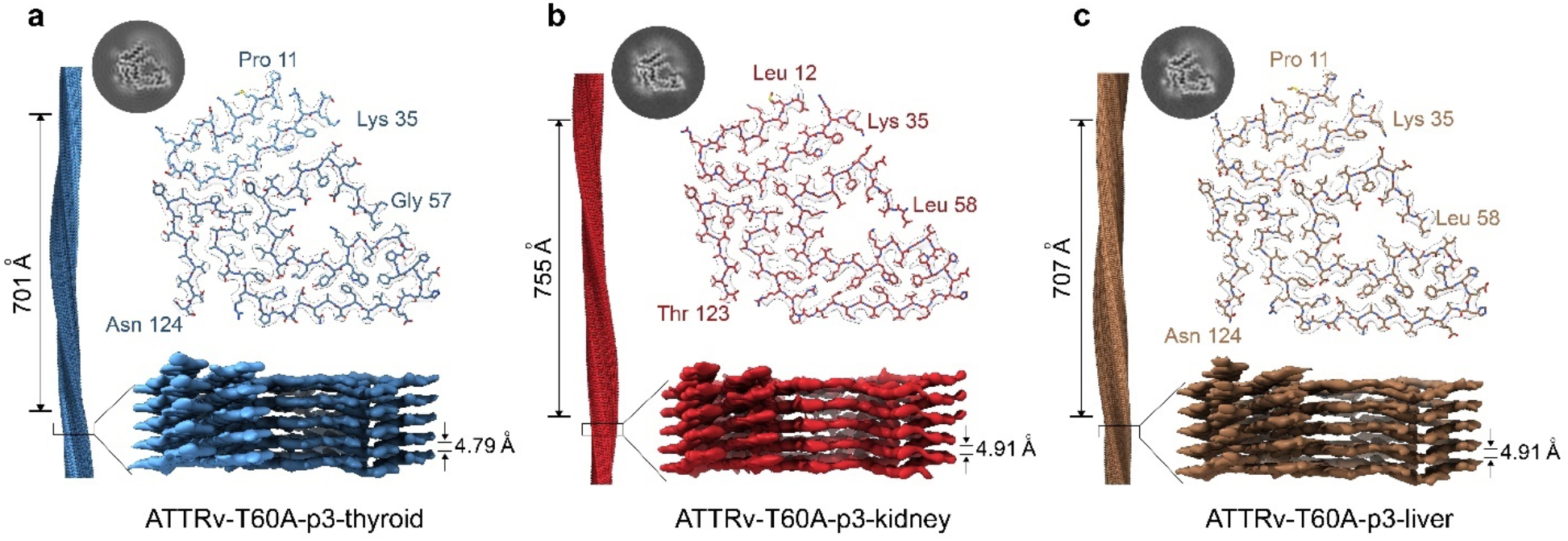
Structure of the fibrils extracted from the thyroid, kidney, and liver of the patient 3. 3D map and models of the fibrils from **a)** thyroid, **b)** kidney, and **c)** liver of ATTRv-T60A fibrils from patient 3 shows the same single morphology for the three of them. Crossover and interlayer distances are indicated. The resulting morphology corresponds to the “closed-gate” fold, in agreement with the structure obtained in the previously analyzed cardiac fibrils.

### The deposition of ATTRv-T60A significantly affects the connective tissue across all examined organs

In line with the symptoms, the organs primarily impacted by ATTR deposition are the heart and nerves, with documented deposition also observed in ligaments and tendons (Sueyoshi et al., 2011). The deposition in other organs is diverse and contingent upon the type of mutation and individual factors (Gertz, 2017), and the mechanisms driving this organ selectivity remain uncertain. The thyroid, studied in this study, is commonly implicated as a key site for ATTR deposition, perhaps due to its integral role in synthesizing thyroxine, the hormone carried by transthyretin (Hofer & Anderson, 1975; Huang et al., 2017). We observe ATTR deposition in the thyroid of patient 3, more specifically in the vascular walls within the thyroid parenchyma and the adipose tissue surrounding the organ **(Figure 2)**.

Contrary, the kidney is rarely affected in ATTR amyloidosis patients (Adams et al., 2021). When it occurs, renal malfunction is caused by cardiorenal syndrome or by direct damage caused by amyloid deposition in the glomeruli, arterioles, and medium vessels (Allinovi et al., 2022; Fenoglio et al., 2022; Lobato et al., 1998). Various mutations have been linked to renal abnormalities, yet the ATTRv-T60A variant has not been specifically addressed (Lobato & Rocha, 2012). In our study, patient 3 underwent combined liver-kidney transplantation, indicating renal dysfunction attributed to ATTR fibril deposition, to prevent the recurrence of nephropathy (Lobato & Rocha, 2012). When analyzing the presence of ATTR deposits in the explanted kidney of patient 3, we observe the presence of amyloid in the adipose tissue surrounding the organ and in the nerve walls within this perirenal fat **(Figure 2)**.

Liver disease has not described as a clinical manifestation of ATTR amyloidosis (Ibrahim et al., 2019), and the rare cases that reported abnormal liver function in ATTR patients were in absence of hepatic amyloid deposits (Bhattacharya et al., 2021). Most histological and immunochemical studies resulted negative for ATTR deposition in the liver and only slight aggregation has been found in the nerves within the parenchyma of the liver (Hofer & Anderson, 1975). In the liver of patient 3, we observe the presence of amyloid deposits in the fat surrounding the falciform ligament of the liver and its vasculature **(Figure 2)**.

### We found subtle differences between ATTRv-T60A fibril extracts

Although all the fibrils examined in this study share the same structure, we found subtle differences in composition. Mass spectrometry data revealed the presence of wild-type and T60A transthyretin in all the fibrils analyzed, consistent with the heterozygosis of the patients **(Figure 4)**. In agreement with previous observations (Liepnieks & Benson, 2007; Liepnieks et al., 2010), the presence of wild-type transthyretin was increased in the fibrils from thyroid and heart of the patient 3, indicating that the deposition of transthyretin continued after liver transplantation.

**Figure 4.**
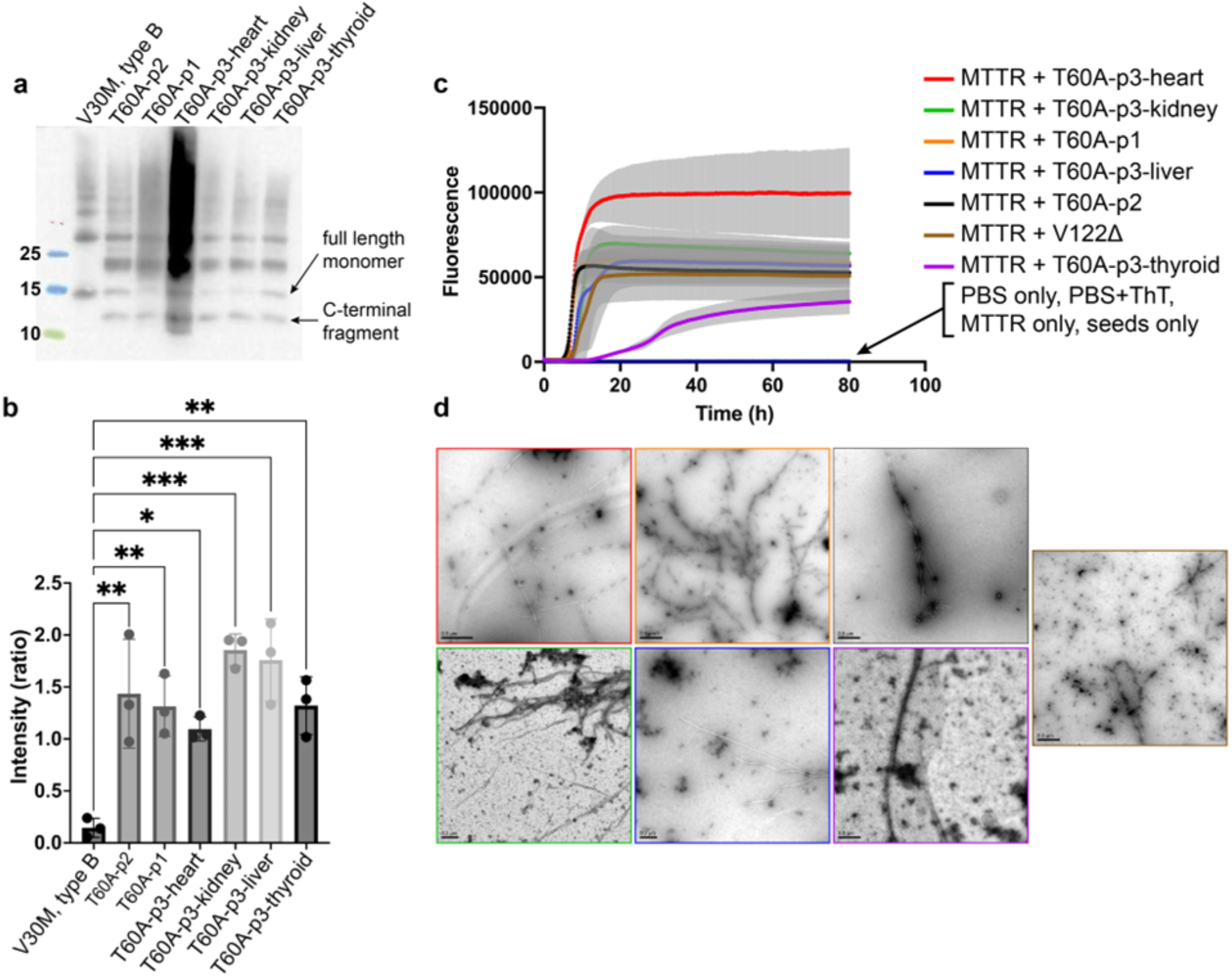
Molecular characterization of the ATTRv-T60A fibrils analyzed in this study. **a)** Western-blot of the different fibrils show the presence of the bands corresponding to the full-length monomeric transthyretin and the c-fragment, according to type A fibrils. **b)** T-test comparison of the ratio C-fragment/full-length does not indicate differences in the cleavage of the ATTRv-T60A fibrils depending on the individual nor the organ. **c)** Cardiac and non-cardiac ATTRv-T60A fibrils seeds the aggregation of MTTR, generating the typical ThT sigmoidal curve, except for that from thyroid. **d)** TEM images after seeding reaction. The presence of fibrils confirmed the success of the seeding reaction.

Additionally, we observe the presence of unknown annular components decorating the renal fibrils from patient 3 **(Supplementary Fig. 3)**. These annular structures of unknown nature could potentially have a role in the aggregation or clearance process. Based on the shape and volume of these structures, we speculate that they correspond to serum amyloid P component (SAP), composed of ten 25.5 kDa subunits non-covalently associated in two pentameric rings, interacting face to face (Emsley et al., 1994). SAP is often found associated with amyloid deposits in diverse diseases (Botto et al., 1997), and its presence was previously reported in renal ATTR deposits (Fenoglio et al., 2022).

### ATTRv-T60A fibrils share a similar proteolytic pattern

The fibrils extracted from all the organs examined here show a similar ratio between C-terminal fragments and full-length transthyretin **(Figure 4 a-b)**. This finding may indicate that the proteolytic cleavage that leads to the formation of C-terminal fragments occurs prior to tissue deposition. An alternative explanation could be the presence of a common protease in all the different deposition sites, or perhaps in the surrounding adipose tissue (Pou et al., 2007; Wu et al., 2023).

### Amyloid seeding of fibrils extracted from the liver of patient 3 could offer valuable insights into clinical observations

In our previous work, we reported the capability of cardiac ATTR fibrils of inducing the aggregation of soluble transthyretin (Saelices et al., 2018). Our present study extends this observation to fibrils extracted from kidney and liver **(Figure 4c-d)**. Several explanted livers from ATTR patients have been widely used for Domino liver transplantation (Marques et al., 2020). In this procedure, the explanted liver carrying the transthyretin mutation, is donated to a patient suffering end stage liver disease or severe metabolic disorder. Given that ATTRv amyloidosis typically takes decades to manifest, this procedure is anticipated to result in a functional liver without leading to significant deposition for the majority of the recipient’s lifetime. However, recipients often develop ATTR-related symptomatology months after the surgery for reasons that are still unknown (Lladó et al., 2010; Stangou et al., 2005). Although we found that the parenchyma of the liver does not have significant amyloid deposits, we found extensive deposition within the Glisson’s capsule and surrounding adipose tissue **(Figure 2)**. These aggregates would be incorporated into the recipient and potentially accelerate the transthyretin aggregation by amyloid seeding (Saelices et al., 2018).

### Closing remarks

Although the widespread distribution of deposits poses a severe challenge, their structural uniformity presents an advantage for drug design targeting ATTR. The insights gained from these cryo-EM studies elucidating the pathological deposits of ATTR provide valuable understanding of transthyretin aggregation mechanisms and precisely identify targets for structure-based diagnostic and therapeutic strategies.

## Materials & Methods

### Tissue samples from patients

We obtained samples of three different patients carrying the mutation T60A, from the laboratory of Dr. Merrill D. Benson at the University of Indiana. We have cardiac tissue of the left ventricle of Patients 1 and 2. We have tissue from the original heart, thyroid, kidney, and liver organs of Patient 3, who underwent a kidney-liver transplantation. The patient with type B ATTR fibrils, carried the mutation V30M and suffered peripheral and autonomic neuropathy. All the tissues were fresh frozen, excepting that from the heart of Patient 3, that was lyophilized. The Office of the Human Research Protection Program exempted the study from Internal Review Board since the specimen was anonymized.

### Extraction of ATTRv-T60A fibrils

Amyloid fibrils from the different tissues were obtained by following a protocol previously described (Schmidt et al., 2019). ∼200 mg of fresh frozen tissue was thawed and minced with a scalpel. This tissue or ∼15 mg of the lyophilized tissue was resuspended into 1 mL Tris-calcium buffer (20 mM Tris, 150 mM NaCl, 2 mM CaCl2, 0.1% NaN3, pH 8.0) and then centrifuged for 5 min. The supernatant was discarded, and the pellet was again resuspended in the same buffer for a new centrifugation. This process was repeated for 4 times. The resulting pellet was resuspended and incubated o/n in 1 mL of 5 mg/mL collagenase Tris-calcium buffer solution, 400 rpm, 37 °C. After 30 minutes of centrifugation (3100×g and 4 °C), the resulting pellet was resuspended in 1 mL Tris–EDTA buffer (20 mM Tris, 140 mM NaCl, 10 mM EDTA, 0.1% NaN3, pH 8.0), for a new 5 min centrifugation. This cycle was repeated 10 times. To obtain the first sample elution, we gently resuspend the pellet with 75-100 μL of ice-cold water (5 mM EDTA, H_2_0, 4°C), centrifuge the sample, and collect the supernatant. We obtained at least 5 elutions per sample. Elutions from non-cardiac tissues were low concentrated and contained additional rest of other tissue/components. We obtained three additional elutions of 50 μL in order to get cleaner samples and more concentrated.

### Immunodot blotting

The presence of transthyretin in the elutions from extractions and the incorporation of MTTR into the seeding reaction was confirmed by dot-blot analysis.

The insoluble fraction of the seeding reaction was obtained by spinning the samples for 1 hour at 13000x*g*.The pellet was resuspended in 6M of Guanidine HCl.

5 μL of the final extraction elutions or the resuspended pellet were blotted onto a nitrocellulose membrane (0.2 μm; Bio-Rad). The transthyretin from the extractions was observed by using anti-transthyretin antibody (1:1000, Genscript) and anti-rabbit secondary antibody (1:1,000; Invitrogen 31460). The presence of MTTR in the pellet was visualized by using HisProbe (Thermo Scientific Fisher) and HRP antibody (1:5000).

### Negative-stained transmission electron microscopy

The elutions obtained from the extractions were visualized by negative-stained transmission electron microscopy in order to examine the quality of the sample. Fibril concentration and purity were considered. Carbon film 300-mesh copper grids (Electron Microscopy Sciences) were glow-discharged and incubated for 2 minutes with the sample. The excess of sample was gently removed with a filter paper to be later incubated with 2 μL of 2% uranyl acetate for 1 minute. The excess of staining solution was removed with the filter paper. Specimens were examined using an FEI Tecnai 12 electron microscope with an accelerating voltage of 120 kV.

### Typing of the ATTR fibrils by western blotting

Around 0.5 μg of sample material was dissolved in tricine SDS sample buffer, heated for 2 min at 85 °C, and loaded on a Novex™ 16% tris-tricine gel system using a Tricine SDS running buffer. The gel content was transferred onto a 0.2 μm nitrocellulose membrane and incubated it with a primary antibody (1:1000) targeting the C-terminal region of the wild-type TTR sequence from GenScript. The secondary antibody was horseradish peroxidase-conjugated goat anti-rabbit IgG (dilution 1:1000, Invitrogen). Transthyretin content was visualized using Promega Chemiluminescent Substrate. The intensity of the bands from C-terminal fragment of the transthyretin and full-length monomer were quantify by ImageJ.

### Amyloid seeding assay

We prepare seeds from the amyloid extracted fibrils (Saelices et al., 2018). For that, the extracts were treated with 1% sodium dodecyl sulfate (SDS) and centrifuged for 1500 x g for 20 min in order to further purify the sample. This purification step was repeated twice, and the soluble fractions were removed. The sample then underwent three washes with sodium phosphate-EDTA (without the addition of 1% SDS) through centrifugation, and was then sonicated in a bath sonicator using cycled of 5 seconds on and 5 sec off for a duration of 10 min, using a minimum amplitude of 30 (Q700 sonicator, Qsonica). We measured the total protein content in the prepared seeds by BCA quantification (Micro BCA™ Protein Assay Kit, Thermo Fisher Scientific). 2% (w/w) seed solution was added to 0.5 mg/mL recombinant MTTR in a final volume of 200 μL, containing 10 μM thioflavin T (ThT) and 1× PBS (pH 7.4). The ThT fluorescence emission was measured at 482 nm with absorption at 440 nm in a FLUOstar Omega (BMG LabTech) microplate reader. The plates (384 Well Optical Btw Plt Polybase Black w/o Lid Non-Treated PS, Thermo Fisher Scientific) were incubated at 37 °C with cycles of 9 min shaking (700 rpm double orbital) and 1 min rest throughout the incubation period. Measurements were taken every 10 min (bottom read) with a manual gain of 1000 fold.

### Mass Spectrometry (MS) assay

0.5 μg of extracted ATTR fibrils were dissolved in a tricine SDS sample buffer, heated at 85 °C for 2 minutes and loaded them in a Novex™ 16% tris-tricine gel system using a Tricine SDS running buffer. The gel was running for 10 minutes and stained with Coomassie dye. The resulting ATTRv-T60A smear was excised from the gel. Samples were digested overnight with trypsin (Pierce) following reduction and alkylation with DTT and iodoacetamide (Sigma–Aldrich). Then, they underwent solid-phase extraction cleanup with an Oasis HLB plate (Waters). The resulting samples were injected onto an Q Exactive HF mass spectrometer coupled to an Ultimate 3000 RSLC-Nano liquid chromatography system. Samples were injected onto a 75 um i.d., 15 cm long EasySpray column (Thermo) and eluted with a gradient from 0-28% buffer B over 90 min. Buffer A contained 2% (v/v) ACN and 0.1% formic acid in water, and buffer B contained 80% (v/v) ACN, 10% (v/v) trifluoroethanol, and 0.1% formic acid in water. The mass spectrometer operated in positive ion mode with a source voltage of 2.5 kV and an ion transfer tube temperature of 300 °C. MS scans were acquired at 120,000 resolution in the Orbitrap and up to 20 MS/MS spectra were obtained in the ion trap for each full spectrum acquired using higher-energy collisional dissociation (HCD) for ions with charges 2-8. Dynamic exclusion was set for 20 sec after an ion was selected for fragmentation.

Raw MS data files were analyzed using Proteome Discoverer v3.0 SP1 (Thermo), with peptide identification performed using a semitryptic search with Sequest HT against the human reviewed protein database from UniProt. Fragment and precursor tolerances of 10 ppm and 0.02 Da were specified, and three missed cleavages were allowed. Carbamidomethylation of Cys was set as a fixed modification, with oxidation of Met set as a variable modification. The false-discovery rate (FDR) cutoff was 1% for all peptides.

To calculate the fraction of WT to Mutant T60A in each tissue, we used the intensity of the peptide 49-70 which is present in all samples in good abundance (PSMs >5) and covers the mutation. Percentage content of WT is calculated by dividing the intensity value for WT by sum of total intensity of WT and Mutant and multiplied by 100 to calculate percentage.

### Cryo-EM sample preparation, data collection, and processing

A 3.5 μL aliquot of the freshly-extracted ATTR fibrils were applied to glow-discharged Quantifoil R 1.2/1.3, 300-mesh, Cu grids. The grid was then blotted with filter paper to remove the excess sample, and plunge frozen into liquid ethane using a Vitrobot Mark IV (FEI). This Cryo-EM sample was screened on the Talos Arctica at the Cryo-Electron Microsopy Facility (CEMF) at University of Texas Southwestern Medical Center (UTSW). Cryo-EM data was collected on a 300 kV Titan Krios microscope (FEI) at the Stanford-SLAC Cryo-EM Center (S^2^C^2^). All movies were recorded using a Falcon4 camera and EPU software (Thermo Fisher Scientific). Raw movies were gain-corrected, aligned and dose-weighted using RELION’s own implemented motion correction program (Zivanov et al., 2020). Contrast transfer function (CTF) estimation was performed using CTFFIND 4.1(Rohou & Grigorieff, 2015). All steps of helical reconstruction, three-dimensional (3D) refinement, and post-process were carried out using RELION 4.0 (Kimanius et al., 2021). Single filaments were picked automatically using Topaz, while double filaments were picked manually (Bepler et al., 2019). Particles were initially extracted using a box size of 768 pixels, then down-scaled to 256 pixels to determine the cross-over distance and the population distribution for each morphology. To further process the data, single filaments and double filaments were extracted in box of 192 pixels and 320 pixels, respectively. Reference-free 2D classification were performed for all datasets to select suitable particles for 3D reconstructions. Fibril helices were assumed to be left-handed. We used an elongated Gaussian blob as an initial reference for 3D classifications of single filaments. For 3D classifications of double filaments, the initial model was generated from 2D class average images using a RELION built-in, script relion_helix_inimodel2d (Scheres, 2020). We performed 3D classifications to select for the best particles leading to the beset reconstructed maps. 3D auto-refinements and CTF refinements were carried out to obtain higher resolution maps. Final maps were post-processed using the recommended standard procedures in RELION, and the overall resolutions were estimated at threshold of 0.143 in the Fourier shell correlation (FSC) curve between two independently refined half-maps.

### Stabilization energy calculation

The stabilization energy per residue was calculated by the sum of the products of the area buried for each atom and the corresponding atomic solvation parameters (M. Sawaya et al., 2021). The overall energy was calculated by the sum of energies of all residues, and assorted colors were assigned to each residue in the solvation energy map.

### Model building

The refined maps of the different morphologies found in this patient were post-processed in RELION before building their models (Terwilliger et al., 2018). We used our previously published model of ATTRv-I84S (PDB code 8TDN) as the template to build the model of XX single fibril. Residue modification, rigid body fit zone, and real space refine zone were performed to obtain the resulting models using COOT v0.9.8.1 (Emsley et al., 2010).

## Acknowledgments

We are very grateful to the patients and families who donate the tissues, without which this study would not have been possible. Special thanks to Dr. Merril Benson and the University of Indiana for providing the material. We want to emphasize our recognition to Dr. Merril Benson because his relevant contributions to the amyloidosis field and his relationship with the affected families for decades. We thank the UTSW Cryo-Electron Microscopy Facility, the UTSW Structural Biology Laboratory, the UTSW Electron Microscopy Core Facility, the national cryo-EM facilities Stanford-SLAC (project CA60) for instrumentation, technical support, and/or data collection. We thank the UTSW Proteomics core for technical assistance with the proteomics experiments.

## Contributions

Conceptualization: L.S., M.C.F.R., B.A.N. Methodology: L.S., M.C.F.R., B.A.N., V.S., S.A. Investigation: M.C.F.R., B.A.N., V.S., S.A., B.E., P.B., L.W., M.P., Y.A., P.S., J.C., A.W. Visualization: L.S., M.C.F.R., B.A.N. Funding acquisition: L.S. Project Administration: L.S. Supervision: L.S. Writing original draft: L.S., M.C.F.R. Writing, review and editing: L.S., M.C.F.R.

## Funding

American Heart Association (Career Development Award 847236)

National Institutes of Health, National Heart, Lung, and Blood Institute (New Innovator Award DP2-HL163810)

Welch Foundation (Research Award I-2121-20220331)

UTSW Endowment (Distinguished Researcher Award from President’s Research Council and start-up funds)

Cryo-EM research was partially supported by the following grants:

National Institutes of Health grant U24GM129547, Department of Energy Office of Science User Facility sponsored by the Office of Biological and Environmental Research

Department of Energy, Laboratory Directed Research and Development program at SLAC National Accelerator Laboratory, under contract DE-AC02-76SF00515

The Electron Microscopy Core Facility at UTSW is supported by the National Institutes of Health (NIH) (1S10OD021685-01A1 and 1S10OD020103-01).

Part of the computational resources were provided by the BioHPC supercomputing facility located in the Lyda Hill Department of Bioinformatics at UTSW. URL: https://portal.biohpc.swmed.edu.

