## Supplementary Figures for "Multi-organ structural homogeneity of amyloid fibrils in ATTRv-T60A amyloidosis patients, revealed by Cryo-EM"

**Supplementary Information**

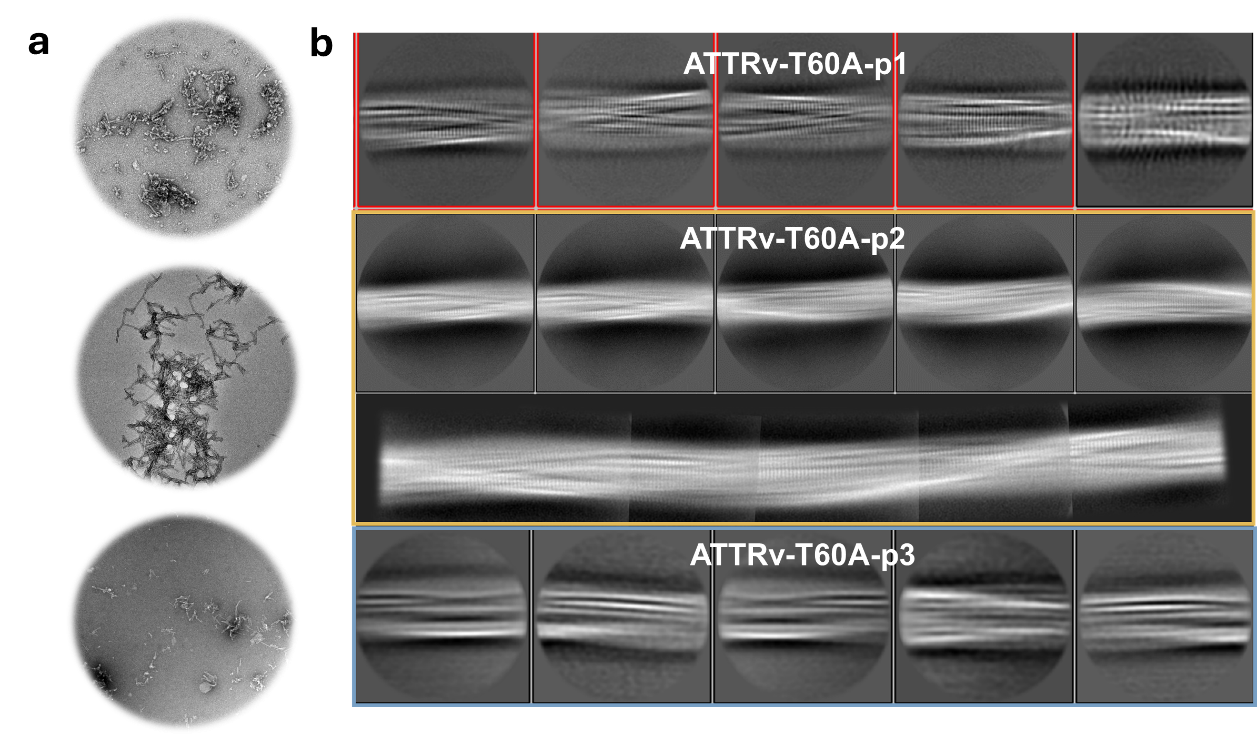

**Supplementary Figure 1. Cardiac ATTRv-T60A fibrils. A)** Electron Microscopy images of the extracted fibrils from the three different patients analyzed in this study. **B)** Representative 2D classes obtained for each patient and 2D fibril reconstruction for ATTRv-T60A-p2.

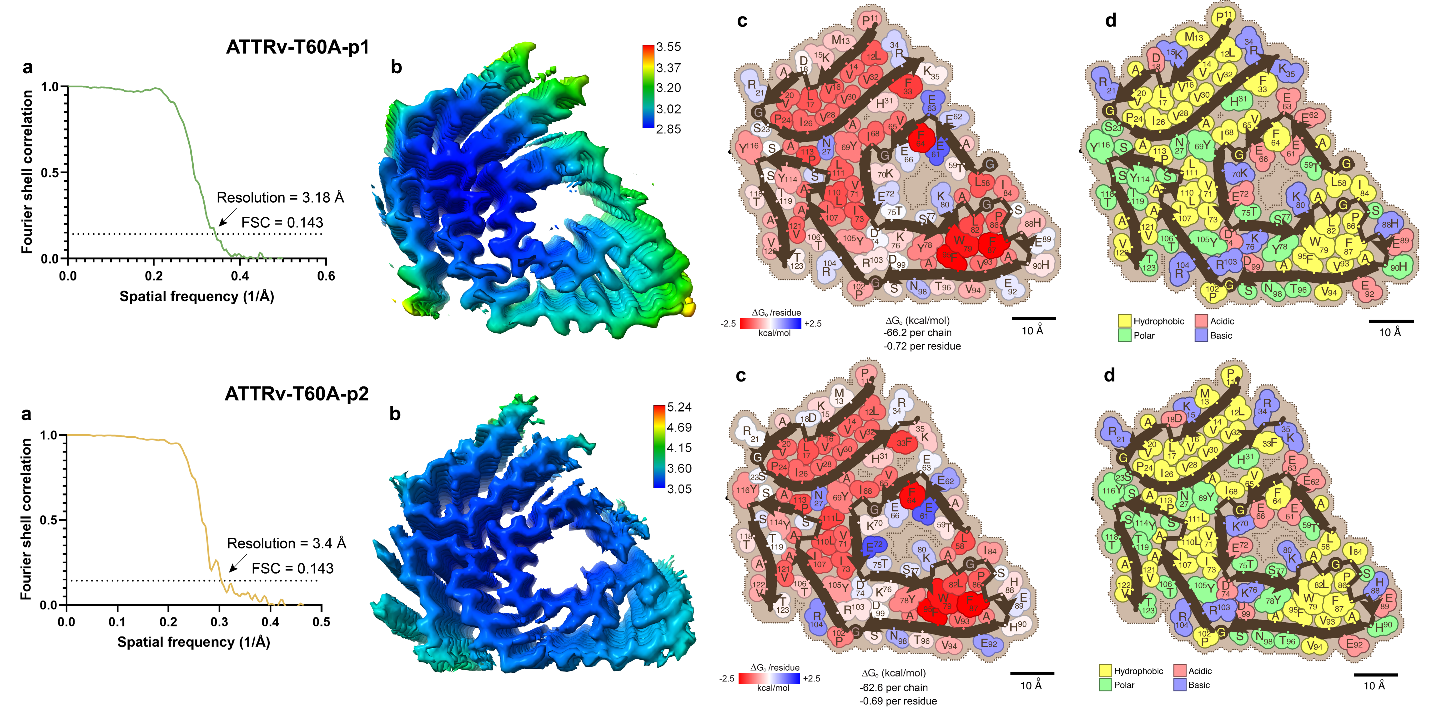

**Supplementary Figure 2. Features of the structures obtained in cardiac ATTRv-T60A fibrils. A)** Final resolution of the density map and **B)** local resolution map of the structure. **C)** The solvation energies shows the three areas (red) that contribute the most to the stability of the fibril. **D)** The residue composition analysis shows how the hydrophobic areas agrees with the most stable regions of the fibril.

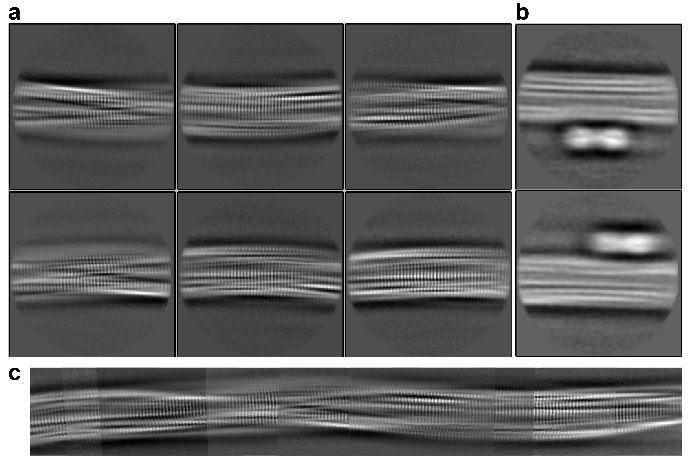

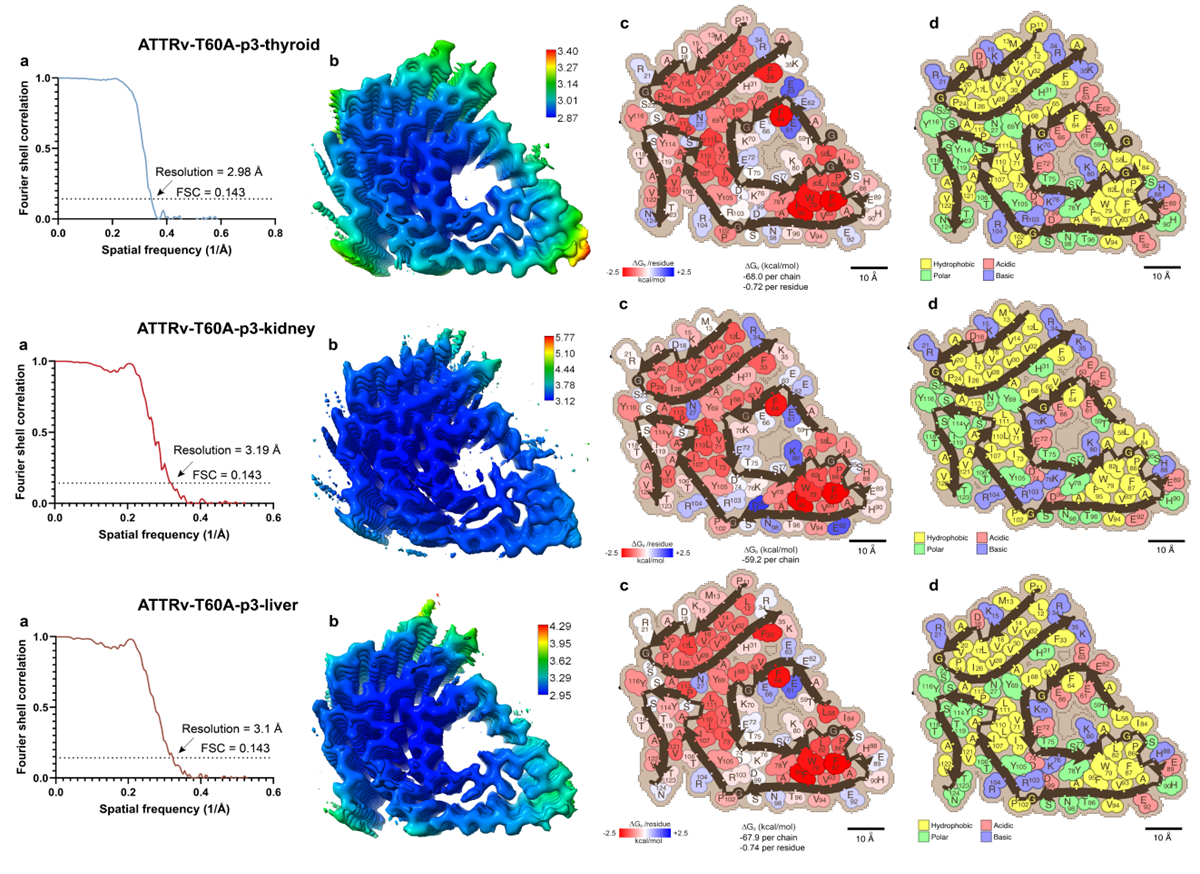
**Supplementary Figure 3. 2D classification in data from kidney. A)** Representative 2D classes in box size of 256 px. Stacked layers, apparent twist, and additional features can be distinguished. **B)** Pairs of annular elements attached to the fibrils. Classes obtained in the first rounds of classification are with low resolution but show additional components that are removed in later iterations. **C)** Stitching of the 2D classes obtained. The 2D view of the fibril morphology can be reconstructed. The classes reconstructing the fibril will be used in the next steps of data processing.

**Supplementary Figure 4. Features of the structures obtained in non-cardiac ATTRv-T60A fibrils. A)** Final resolution of the density map and **B)** local resolution map in the structure. **C)** The solvation energies and the **D)** residue composition analysis shows similar values and features with those obtained in cardiac fibrils.

**Supplementary Table 1.** Data collection and model refinement statistics.

|  | **T60A-p1** | **T60A-p2** | **T60A-p3-h** | **T60A-p3-t** | **T60A-p3-k** | **T60A-p3-l** |
| --- | --- | --- | --- | --- | --- | --- |
| **Microscope (Model)** | Titan Krios | Titan Krios | Titan Krios | Titan Krios | Titan Krios | Titan Krios |
| **Acceleration Voltage (kV)** | 300 | 300 | 300 | 300 | 300 | 300 |
| **Detector** | Falcon 4i | K3 | K3 | K3 | Falcon 4i | Falcon 4i |
| **Software** | EPU 3.5 | SerialEM 3.8 | SerialEM 3.8 | SerialEM 3.8 | EPU 3.5 | EPU 3.5 |
| **Magnification** | 130,000x | 81,000 | 105,000x | 105,000x | 130,000 | 130,000 |
| **Pixel size at detector (Å/px)** | 0.946 | 0.533 | 0.834 | 0.834 | 0.946 | 0.954 |
| **Defocus range (µm)** | -0.8 to -1.9 | -1.4 to -2.2 | -0.9 to -2.2 | -0.9 to -2.2 | -0.9 to -2.1 | -1.2 to -2.8 |
| **Total dose (e)** | 40 | 42 | 50 | 50 | 40 | 40 |
| **Exposure time (sec)** | 4.8 | 2.2 | 1.80 | 1.80 | 5.29 | 5.93 |
| **Number of movie frames** | 40 | 57 | 50 | 50 | 40 | 40 |
| **Usable micrographs** | 8,293 | 3735 | 6,064 | 6,236 | 6,699 | 6,691 |
| **Total extracted segments** | 2,513,579 | 261,754 | 207,988 | 675,457 | 1,271,167 | 727,422 |
| **Box size** | 256 | 256 | 256 | 256 | 256 | 256 |
| **Number of segments after 2D** | 757,654 | 226,710 | 67016 | 479,155 | 464,905 | 461,912 |
| **Number of straight segments** | 23,165 | 18,467 | 3168 | 0 | 16,697 | 4,083 |
| **Number of segments after 3D** | 104,700 | 55,692 | 49669 | 133,439 | 57,895 | 30,791 |
| **Symmetry imposed** | C1 | C1 | C1 | C1 | C1 | C1 |
| **Helical rise (Å)** | 4.90 | 4.78 | n/a | 4.79 | 4.91 | 4.91 |
| **Helical twist (°)** | -1.26 | -1.27 | n/a | -1.23 | -1.17 | -1.25 |
| **Crossover length (Å)** | 700 | 677 | ~690 | 701 | 755 | 707 |
| **B factor** | -121.7 | -83.5 | -184 | -70.5 | -134.9 | -112.9 |
| **Map resolution (Å; FSC=0.143)** | 3.18 | 3.4 | 4.64 | 2.98 | 3.19 | 3.10 |
| **Map resolution (Å; FSC=0.5)** |  |  |  |  |  |  |
| **Non-hydrogen atoms** | 3,580 | 3,560 | n/a | 3,645 | 3,525 | 3,600 |
| **Protein residues** | 460 | 455 | n/a | 470 | 450 | 460 |
| **Number of chains** | 5 | 5 | n/a | 5 | 5 | 5 |
| **Water/ligands** | 0 | 0 | n/a | 0 | 0 | 0 |
| **MolProbity score** | 1.49 | 2.25 | n/a | 1.93 | 1.86 | 1.80 |
| **Clash score** | 3.91 | 18.53 | n/a | 12.77 | 10.50 | 9.03 |
| **Rotamer outliers (%)** | 0 | 0 | n/a | 0 | 0 | 0 |
| **R.M.S deviations bonds (Å)** | 0.003 | 0.004 | n/a | 0.002 | 0.004 | 0.005 |
| **R.M.S deviations angle (°)** | 0.655 | 0.821 | n/a | 0.584 | 0.592 | 0.874 |
| **Ramachandran plot** (5)  Favored  Allowed  Outliers | 95.45  4.55  0 | 91.95  8.05  0 | n/a | 95.56  4.44  0 | 95.35  4.65  0 | 95.45  4.55  0 |
| **CaBLAM outliers (%)** | 3.57 | 2.41 | n/a | 2.33 | 2.44 | 2.38 |
| **Model vs Data** | 0.84 | 0.70 | n/a | 0.86 | 0.79 | 0.81 |
